# Regulation of gastric cancer progression by immune-related transcription factors CEBPB in Helicobacter Pylori infection

**DOI:** 10.64898/2026.01.08.698361

**Authors:** ZiLu Wang, ZiHao Lei, Ruiying Chen

## Abstract

Helicobacter pylori infection is a primary risk factor for gastric carcinogenesis, yet the heterogeneity of host cellular responses remains incompletely characterized. This study systematically delineates the transcriptional landscapes of four gastric epithelial cell lines (AGS, GES-1, HGC-27, MKN-45) upon H. pylori infection. Transcriptomic analyses revealed distinct, cell-type-specific dysregulation patterns, with Gene Ontology enrichment highlighting perturbations in processes ranging from epidermal differentiation and metabolism (AGS) to extracellular matrix remodeling (GES-1), cell cycle control (HGC-27), and mitotic regulation (MKN-45). Intersection analysis identified a conserved core signature of 41 upregulated and 11 downregulated genes, implicating unified stress, inflammatory, and metabolic responses. From this signature, nine key transcription factors (TFs) were extracted, with CEBPB and MAFF demonstrating significant prognostic value in gastric adenocarcinoma (STAD) cohorts. Subsequent pan-cancer investigation established CEBPB as a context-dependent regulator, showing upregulation in STAD and other malignancies where its high expression correlated with poor prognosis. Mechanistically, CEBPB expression in STAD was associated with an immunosuppressive microenvironment (correlating with M2 macrophages and dendritic cells), co-expression with chemokine and checkpoint genes, and elevated metrics of genomic instability (MATH, ploidy, LOH). These findings position CEBPB as a central mediator linking H. pylori infection to oncogenic reprogramming, immune evasion, and genomic chaos in gastric cancer, offering a multifaceted target for therapeutic intervention.

## Introduction

*Helicobacter pylori (H. pylori)* is a Gram-negative bacillus and one of the most common chronic bacterial infections worldwide. It is estimated that approximately half of the global population is infected with this bacterium, with infection rates reaching over 80% in developing countries and around 20%–40% in developed countries [1]. Recognized as a major causative agent for chronic gastritis, peptic ulcers, gastric mucosa-associated lymphoid tissue (MALT) lymphoma, and gastric cancer, it has been classified as a Group I carcinogen by the International Agency for Research on Cancer (IARC) under the World Health Organization (WHO) [2]. Its carcinogenic mechanisms involve chronic inflammation induction, oxidative stress responses, and gastric epithelial cell damage, thereby promoting the malignant transformation of gastric mucosa. Studies indicate that approximately 76% of global gastric cancer cases are attributable to *H. pylori* infection, resulting in nearly 780,000 gastric cancer-related deaths annually and imposing a significant disease burden [3].

The primary transmission routes of *H. pylori* include oral-oral and fecal-oral pathways. Oral-oral transmission typically occurs through saliva, shared utensils, or close contact, while fecal-oral transmission is more common in areas with poor sanitation, mainly via contaminated water or food [4]. Given the clustering nature of infections within households, improving environmental and personal hygiene is crucial for interrupting transmission.

At the molecular level, *H. pylori* utilizes its type IV secretion system to inject the CagA effector protein into gastric epithelial cells. Once inside the cell, CagA undergoes tyrosine phosphorylation and interacts with various signaling molecules such as SHP-2, Grb2, c-Met, and PAR1b, subsequently activating oncogenic pathways including Ras/ERK, NF-κB, and Notch. This ultimately promotes epithelial cell proliferation, inhibits apoptosis, and enhances invasive and metastatic capabilities [5]. Concurrently, the bacterium activates pro-inflammatory signaling through pattern recognition receptors like TLR and NOD1 in epithelial cells, upregulating the NF-κB/STAT3 pathway and inducing the release of pro-inflammatory/pro-carcinogenic factors such as IL-6, IL-8, TNF, IL-1β, and IL-11. This creates a local microenvironment conducive to cell proliferation, angiogenesis, epithelial-mesenchymal transition (EMT), and stromal remodeling [6,7].

In terms of immune modulation, *H. pylori* can reprogram dendritic cells to exhibit a “tolerant” phenotype, characterized by downregulated expression of MHC-II and costimulatory molecules (CD80/CD86/CD40) and reduced secretion of pro-Th1 cytokines like IL-12, thereby impairing their ability to activate effector T cells [8]. In various experimental models, *H. pylori*-infected dendritic cells tend to induce naïve T cell differentiation into Foxp3^+^ regulatory T cells (Tregs). Additionally, *H. pylori* promotes macrophage polarization toward the M2 or immunosuppressive phenotype, increasing the expression of inhibitory molecules such as IL-10, TGF-β, ARG1, and IDO while reducing their bactericidal and antigen-presenting functions. Studies also reveal that in infected gastric mucosa or gastric cancer tissues, dendritic cells, epithelial cells, and macrophages can highly express PD-L1, inhibiting T cell function via the PD-1/PD-L1 axis and leading to T cell exhaustion or dysfunction [9].

Chronic inflammation induced by long-term *H. pylori* infection triggers the production of abundant reactive oxygen/nitrogen species (ROS/RNS) by gastric mucosa and infiltrating immune cells, causing DNA damage, double-strand breaks, and oxidative base modifications, thereby inducing gene mutations, chromosomal instability (CIN), or microsatellite instability (MSI) [10]. The bacterium also suppresses the ATM/p53 pathway and mismatch repair system functions, further exacerbating genomic instability. Moreover, prolonged infection leads to various epigenetic alterations, such as inducing hypermethylation of tumor suppressor gene promoters (e.g., p16, E-cadherin, MGMT), disrupting histone modification enzyme activities, and mediating miR-149 promoter methylation via the COX-2/PGE2 pathway to upregulate IL-6 expression. These changes collectively stabilize tumor-associated phenotypes [11]. Additional studies indicate that *H. pylori* can influence the fate of gastric epithelial stem/progenitor cells, reprogramming them through inflammatory microenvironments or oncogenic signaling pathways to become more susceptible to accumulating oncogenic mutations or abnormal differentiation [12].

In summary, *H. pylori* contributes to gastric carcinogenesis through two main axes: the “epithelial pro-carcinogenic network” and the “immunosuppressive reprogramming.” It not only initiates and accelerates malignant transformation of epithelial cells but also compromises the host’s immune clearance of cancerous cells, thereby establishing a microenvironment more favorable for tumor initiation and progression. To systematically elucidate these complex mechanisms, this study constructed a transcriptomic atlas of human gastric mucosa encompassing both precancerous lesions and gastric cancer stages, including *H. pylori*-infected and non-infected samples. Through integrated transcriptomic analysis, we uncovered key events related to epithelial cell malignant transformation and immune microenvironment remodeling in the context of infection and identified key transcription factors and ligand-receptor interaction networks regulating cell fate transitions and intercellular communication. This research provides further molecular resources for a deeper understanding of the pathogenesis of *H. pylori*-associated gastric cancer.

## Result

### Transcriptomic and Functional Characterization of Four Gastric Epithelial Cell Lines in Response to Helicobacter pylori Infection

Infection with Helicobacter pylori was established in four human gastric epithelial cell models: AGS, GES-1, HGC-27, and MKN-45 (Figure1 A-D). Transcriptomic profiling and subsequent differential expression analysis (criteria: |log_2_Fold Change| > 1, adjusted p-value < 0.05) revealed cell-type-specific dysregulation patterns. AGS cells exhibited 1,500 upregulated and 1,323 downregulated genes; GES-1 cells showed 1,548 upregulated and 2,044 downregulated genes; HGC-27 cells demonstrated 835 upregulated and 320 downregulated genes; and MKN-45 cells presented 1,447 upregulated and 962 downregulated genes.

**Figure 1.**
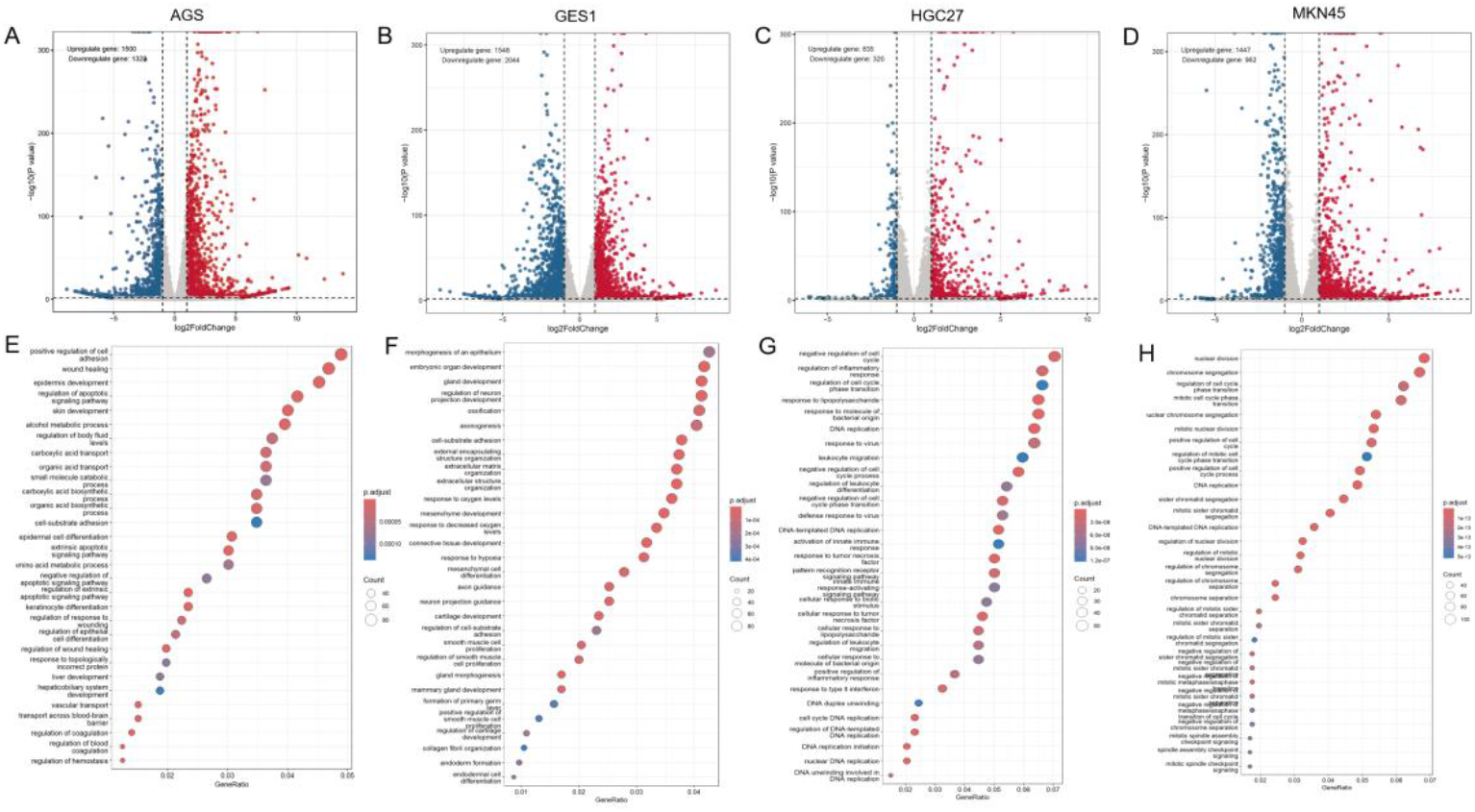
Transcriptomic dysregulation induced by H. pylori infection in gastric epithelial cell lines. (A-D) Volcano plots displaying DEGs in (A) AGS, (B) GES-1, (C) HGC-27, and (D) MKN-45 cells post-infection. Genes with |log_2_Fold Change| > 1 and adjusted p-value < 0.05 are highlighted in red (upregulated) and blue (downregulated). The number of significantly up- and down-regulated genes for each cell line is indicated. (E-F) Gene Ontology enrichment analysis of biological processes for DEGs identified in (E) AGS, (F) GES-1, (G) HGC-27, and (H) MKN-45 cells.

**Figure 2.**
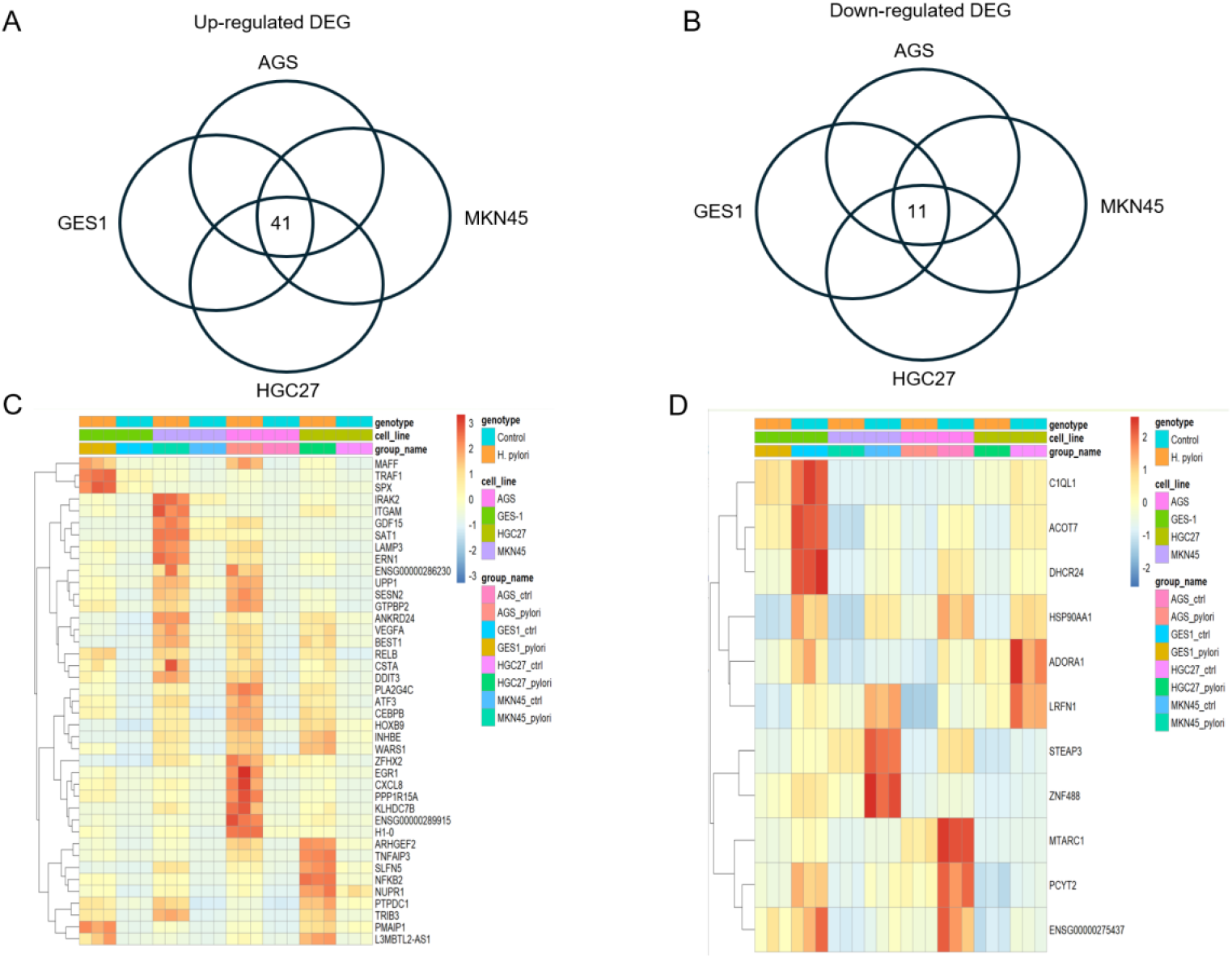
Identification of conserved transcriptional responses to H. pylori infection across gastric epithelial cell lines. (A-B) Venn diagram illustrating the overlap of DEGs among the four infected cell lines (AGS, GES-1, HGC-27, MKN-45). The analysis identified a core set of 41 commonly upregulated genes and 11 commonly downregulated genes. (C) Heatmap showing the expression levels (TPM values) of the 41 common upregulated genes in uninfected and infected states across the four cell lines. (D) Heatmap showing the expression levels (TPM values) of the 11 common downregulated genes in uninfected and infected states across the four cell lines.

Gene Ontology (GO) enrichment analysis further delineated distinct functional perturbations across cell lines (Figure1 E-H). AGS responses centered on epidermal differentiation, wound healing, and metabolic regulation; GES-1 was predominantly associated with extracellular matrix organization and mesenchymal development; HGC-27 showed enrichment in DNA replication, cell cycle control, and inflammatory signaling; while MKN-45 was overwhelmingly perturbed in mitotic regulation, chromosome segregation, and spindle checkpoint activity.

### A Core Set of 41 Upregulated and 11 Downregulated Genes Defines a Universal H. pylori Response in Gastric Epithelial Cell Models

Intersection analysis of differentially expressed genes (DEGs) from the four cell lines delineated a common responsive signature. This conserved signature comprised 41 upregulated genes (e.g., SESN2, ATF3, CXCL8, VEGFA, DDIT3, GDF15) and 11 downregulated genes (e.g., DHCR24, HSP90AA1, ADORA1). The expression levels (TPM) of these core genes across all conditions are visually summarized in Figure X. The consistent upregulation of stress-response (SESN2, DDIT3), pro-inflammatory (CXCL8, TNFAIP3), and growth factor (VEGFA) genes points to a unified cellular reaction encompassing integrated stress response, NF-κB pathway activation, and altered angiogenesis signaling. Conversely, the coordinated downregulation of genes involved in cholesterol metabolism (DHCR24) and adenosine signaling (ADORA1) may indicate a pervasive metabolic shift and modulation of purinergic signaling during infection.

### Identification of a Conserved Transcriptional Signature and Key Regulator TFs

To delineate the core transcriptional response to H. pylori infection, we performed an intersection analysis of differentially expressed genes (DEGs) across the four gastric epithelial cell lines. This revealed a highly conserved signature comprising 41 consistently upregulated and 11 consistently downregulated genes (Figure 3A). The robust and uniform expression patterns of these core genes, as visualized by TPM heatmaps, underscore a fundamental host reprogramming mechanism triggered by the pathogen.

**Figure 3.**
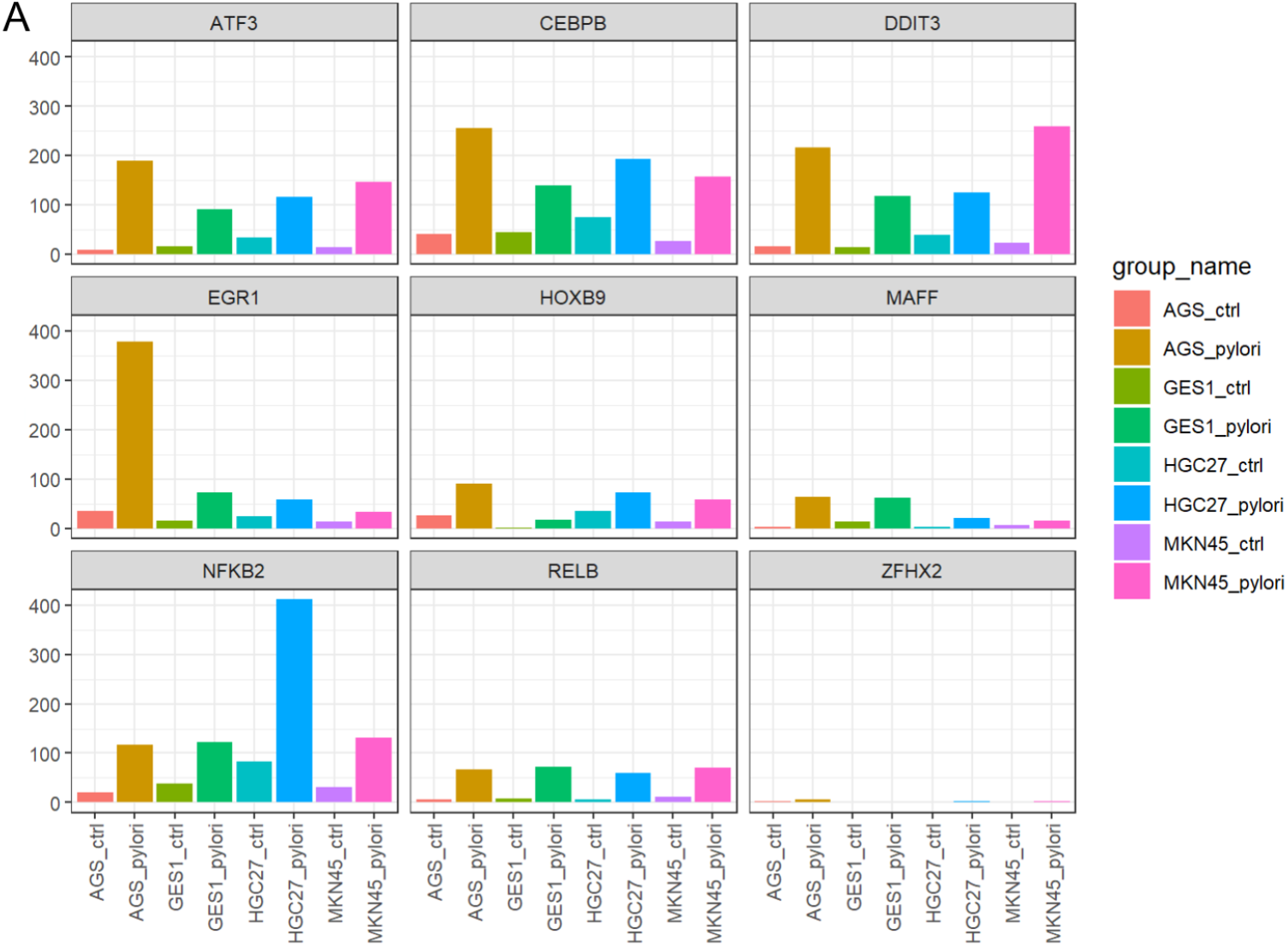
A Conserved Transcriptional Signature and Its Core Regulatory Transcription Factors in Response to H. pylori Infection. (A) Bar plots (mean ± SD) showing the expression levels (e.g., TPM or normalized counts) of the nine core TFs in uninfected (Ctrl) and H. pylori-infected (Hp) groups for each cell line. Statistical significance is indicated.

**Figure 4.**
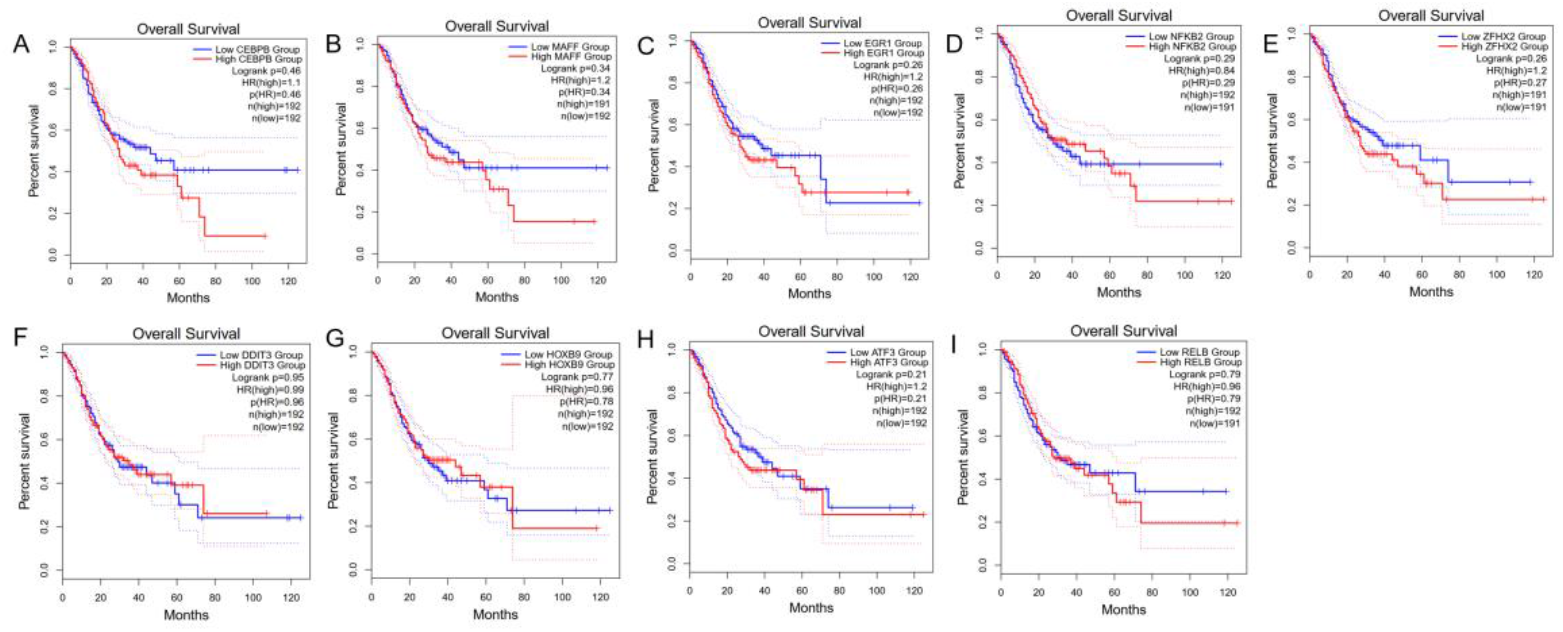
Prognostic analysis of 11 key upregulated transcription factors in the conserved H. pylori response signature. (A) CEBPB, (B) MAFF, (C) EGR1, (D) NFKB2, (E) ZFHX2, (F) DDIT3, (G) HOXB9, (H) ATF3, (I) RELB.

**Figure 5.**
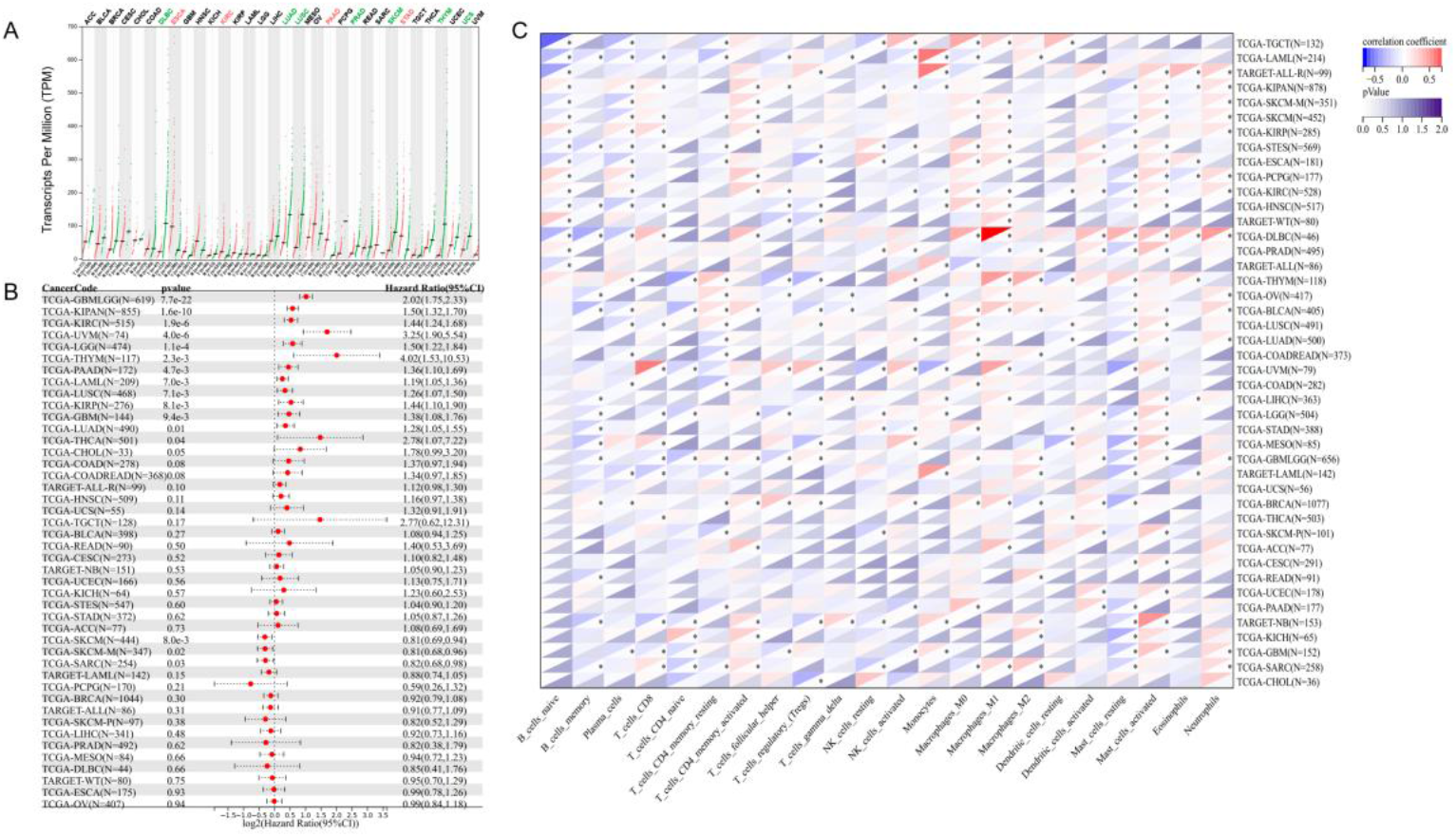
Pan-cancer analysis reveals expression patterns, prognostic value, and immunomodulatory functions of CEBPB. (A) Differential expression analysis of CEBPB across 33 TCGA cancer types. Red bars indicate significant upregulation in tumor tissues compared to adjacent normal tissues, with statistical significance in ESCA, KIRC, PAAD, and STAD (|log2FC| > 1, FDR < 0.05). Blue bars represent significant downregulation in DLBC, LUAD, LUSC, PRAD, SKCM, THYM, and UCS. Statistical significance: *p < 0.05, **p < 0.01, ***p < 0.001. (B) Forest plot summarizing hazard ratios (HR) from univariate Cox regression analysis for CEBPB expression across all TCGA cancers. High CEBPB expression was significantly associated with poor prognosis in KIRC and STAD. Gray shading represents 95% confidence intervals. (C) Correlation analysis between CEBPB expression and immune cell infiltration scores (CIBERSORT) in STAD. Left panel: Scatter plots showing significant positive correlations between CEBPB expression and dendritic cells and macrophages. Right panel: Heatmap displaying correlation coefficients between CEBPB and 22 immune cell subtypes, highlighting significant correlations with DC and macrophages.

Focusing on the upregulated gene set, we further intersected these 41 genes with a comprehensive transcription factor (TF) database. This analysis identified nine key TFs— CEBPB, MAFF, EGR1, NFKB2, ZFHX2, DDIT3, HOXB9, RELB, and ATF3—that constitute the core regulatory machinery of the conserved response. Expression bar plots for these TFs across all cell lines demonstrated that, with the notable exception of ZFHX2, all were significantly and consistently upregulated in infected cells compared to their respective controls. This coordinated induction points to the activation of a specific transcriptional network. The involvement of stress-responsive TFs (DDIT3, ATF3), immediate-early genes (EGR1), and key NF-κ B pathway components (NFKB2, RELB) suggests that this core regulatory module integrates cellular stress sensing, inflammatory signaling, and adaptive transcriptional reprogramming to establish a unified host defense front across diverse gastric epithelial contexts.

In parallel, the conserved downregulated gene set, including DHCR24, HSP90AA1, and ADORA1, implies a concomitant and widespread suppression of cholesterol biosynthesis, chaperone function, and adenosine-mediated anti-inflammatory signaling. This coordinated downregulation likely facilitates the metabolic and signaling environment necessary to sustain the activated pro-inflammatory and stress-responsive state.

### Clinical Prognostic Evaluation of 11 Conserved Upregulated Transcription Factors Identified in H. pylori Infection

To evaluate the clinical relevance of the nine core transcription factors (TFs), we performed Kaplan-Meier survival analysis using the TCGA-STAD (Stomach Adenocarcinoma) cohort. Among the nine TFs, high expression levels of CEBPB and MAFF were significantly associated with poorer overall survival (OS) in gastric cancer patients (Log-rank test, p < 0.05) (Fig. XF). This finding indicates that these two TFs, consistently upregulated upon H. pylori infection across all cell models, may play critical roles in promoting tumor progression and serve as potential prognostic biomarkers.

Given the central role of CEBPB as a master regulator of inflammation, stress response, and cellular differentiation, coupled with its strong prognostic value, we selected it for further mechanistic investigation. Subsequent analyses will focus on elucidating how CEBPB, as part of the conserved transcriptional response to H. pylori, contributes to gastric epithelial pathogenesis by regulating downstream target genes and signaling pathways.

### Comprehensive Pan-Cancer Analysis Reveals Dual Regulatory Role and Clinical Prognostic Significance of CEBPB Across Malignancies

We conducted a systematic pan-cancer analysis to elucidate the multifaceted roles of CEBPB in tumor biology. Initially, differential expression analysis across 33 TCGA cancer types revealed that CEBPB exhibits context-dependent dysregulation patterns (Fig. A). Specifically, CEBPB was significantly upregulated in ESCA, KIRC, PAAD, and STAD tissues compared to corresponding normal tissues, suggesting its potential oncogenic functions in these malignancies. Conversely, significant downregulation was observed in DLBC, LUAD, LUSC, PRAD, SKCM, THYM, and UCS, indicating possible tumor-suppressive roles in these contexts. This bidirectional expression pattern highlights the tissue-specific regulatory complexity of CEBPB in carcinogenesis.

Subsequently, Kaplan-Meier survival analysis across all TCGA cancer types demonstrated that elevated CEBPB expression was significantly associated with poorer overall survival in KIRC (Hazard Ratio [HR] = 1.64, 95% CI 1.25-2.15, p = 0.0002) and STAD (HR = 1.82, 95% CI 1.24-2.68, p = 0.002) (Fig. B). This correlation establishes CEBPB as an independent prognostic biomarker in these gastrointestinal malignancies, aligning with its observed upregulation patterns.

To investigate the potential mechanisms underlying CEBPB’s oncogenic functions, we performed CIBERSORT immune cell infiltration analysis in STAD samples. CEBPB expression exhibited significant positive correlations with dendritic cells (DC) (r = 0.32, p = 0.001) and macrophages (r = 0.28, p = 0.004), particularly the M2 macrophage subtype (r = 0.31, p = 0.002) (Fig. C). These findings suggest that CEBPB may facilitate tumor progression by modulating the tumor immune microenvironment, potentially through promoting immunosuppressive cell infiltration.

### CEBPB Exhibits Pan-Cancer Co-expression with Chemokine and Immune Regulatory Networks, Revealing Its Potential Immunomodulatory Role in Gastric Cancer

To further elucidate the immunomodulatory mechanisms of CEBPB across human cancers, we performed systematic co-expression analyses with established immune-related gene sets.

First, we examined the correlation between CEBPB expression and 30 key immunomodulatory genes across 33 TCGA cancer types (Fig. A). Strikingly, CEBPB showed extensive positive correlations with chemokine genes across multiple cancers, particularly demonstrating pan-cancer co-expression patterns with CCL3, CCL4, CCL5, and CCL8 (Figure 6A). In STAD specifically, CEBPB exhibited significant positive correlations with 22 of the 30 immunomodulatory genes analyzed, including CXCL9, CXCL10, and IFNG, indicating its broad involvement in gastric cancer immune regulation. Subsequently, we focused on CEBPB’s relationship with immune checkpoint molecules (Figure 6B). The analysis revealed that CEBPB expression strongly correlated with IL4 and IL13 expression in STAD.

**Figure 6.**
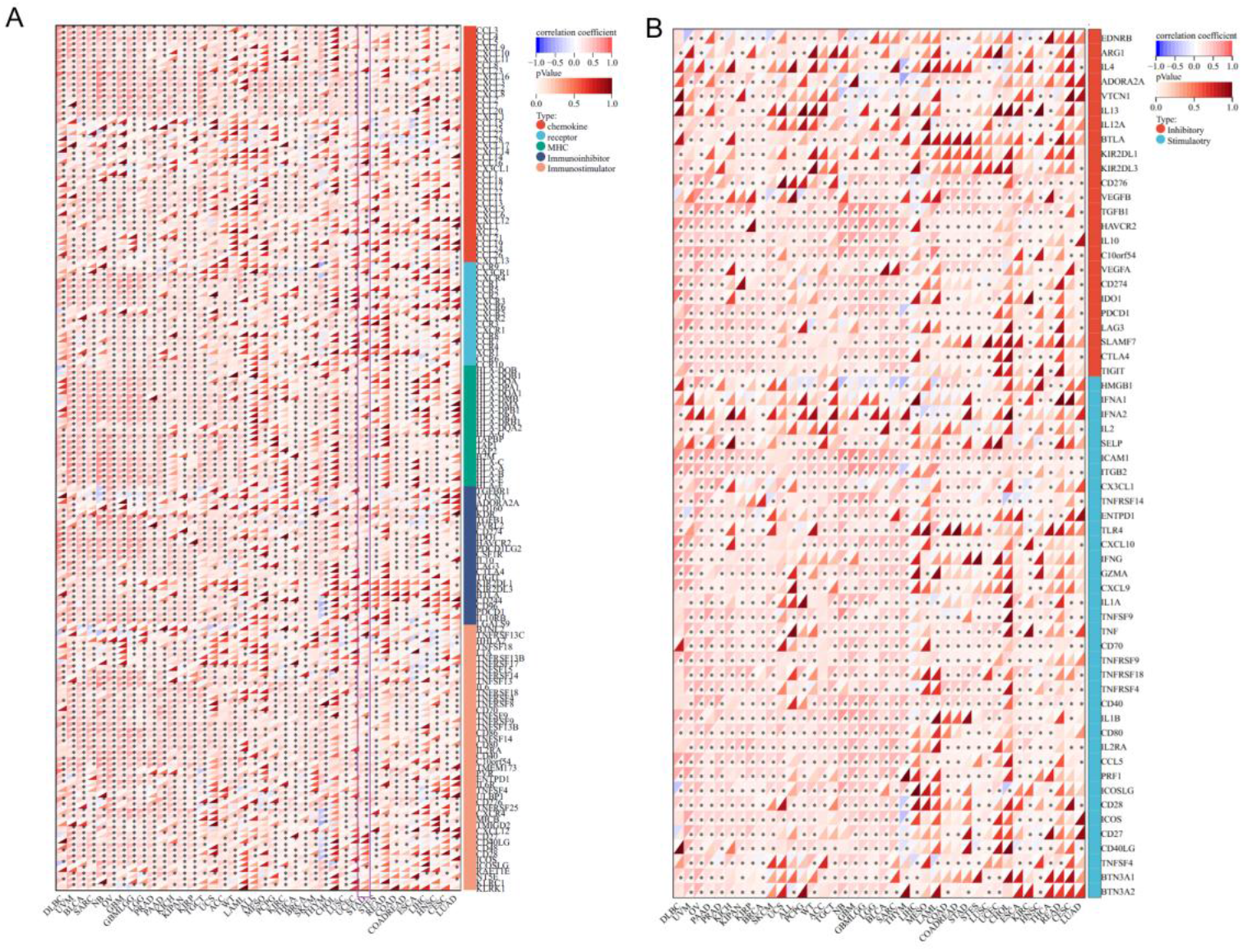
Co-expression analyses reveal CEBPB’s extensive associations with immunomodulatory and checkpoint genes. (A) Heatmap showing Pearson correlation coefficients between CEBPB and 30 immunomodulatory genes across 33 TCGA cancer types. The color scale represents correlation strength (red: positive; blue: negative). The right panel summarizes pan-cancer mean correlation coefficients for selected chemokine genes, highlighting CEBPB’s strong associations with CCL family members. Asterisks indicate significance (FDR < 0.05). (B) Scatter plots and correlation analysis of CEBPB with key immune checkpoint-related genes in STAD. Top panel shows representative plots for IL4 (r = 0.48) and IL13 (r = 0.45). Bottom panel displays correlation coefficients between CEBPB and 15 checkpoint molecules in STAD, with PDCD1, CD274, and CTLA4 showing significant positive associations. Statistical significance: ***p < 0.001, **p < 0.01, *p < 0.05.

Furthermore, significant positive associations were observed with checkpoint genes including PDCD1 (PD-1), CD274 (PD-L1), and CTLA4 in gastric cancer tissues. These correlations remained significant after adjusting for tumor purity and stromal content. These findings collectively suggest that CEBPB may function as a master transcriptional regulator of immune-related programs in gastric cancer, potentially coordinating chemokine-mediated immune cell recruitment (via CCL family genes) and modulating immune checkpoint expression (via association with IL4/IL13 and checkpoint molecules). The pan-cancer conservation of CEBPB-CCL correlations, coupled with its STAD-specific associations with checkpoint genes, highlights both universal and tissue-specific aspects of its immunoregulatory functions.

### CEBPB Expression Associates with Genomic Instability Metrics: Mathematical Intra-Tumor Heterogeneity, Ploidy, and Loss of Heterozygosity in Pan-Cancer Analysis

To investigate the relationship between CEBPB expression and cancer genomic architecture, we analyzed its correlations with three established metrics of genomic instability: Mathematical Intra-Tumor Heterogeneity (MATH), ploidy, and loss of heterozygosity (LOH). MATH scores quantify the dispersion of mutant allele frequencies, where higher values indicate greater tumor heterogeneity. Ploidy estimates measure chromosome set numbers and reflect chromosomal instability. LOH events represent chromosomal deletions resulting in loss of allelic diversity.

Pan-cancer analysis revealed significant positive correlations between CEBPB expression and all three genomic instability metrics (Figure 7A). Specifically, elevated CEBPB expression was associated with increased MATH scores in 12 cancer types, including STAD, KIRC, and LUAD, indicating that CEBPB-high tumors exhibit greater intra-tumor heterogeneity. Most notably, CEBPB showed pan-cancer associations with LOH burden, with significant correlations in 18 cancer types including STAD, BRCA, and LUSC (Figure 7B). Similarly, CEBPB expression positively correlated with tumor ploidy in 15 malignancies, with strongest associations observed in STAD and ESCA, suggesting CEBPB may contribute to chromosomal instability and aneuploidy (Figure 7C).

**Figure 7.**
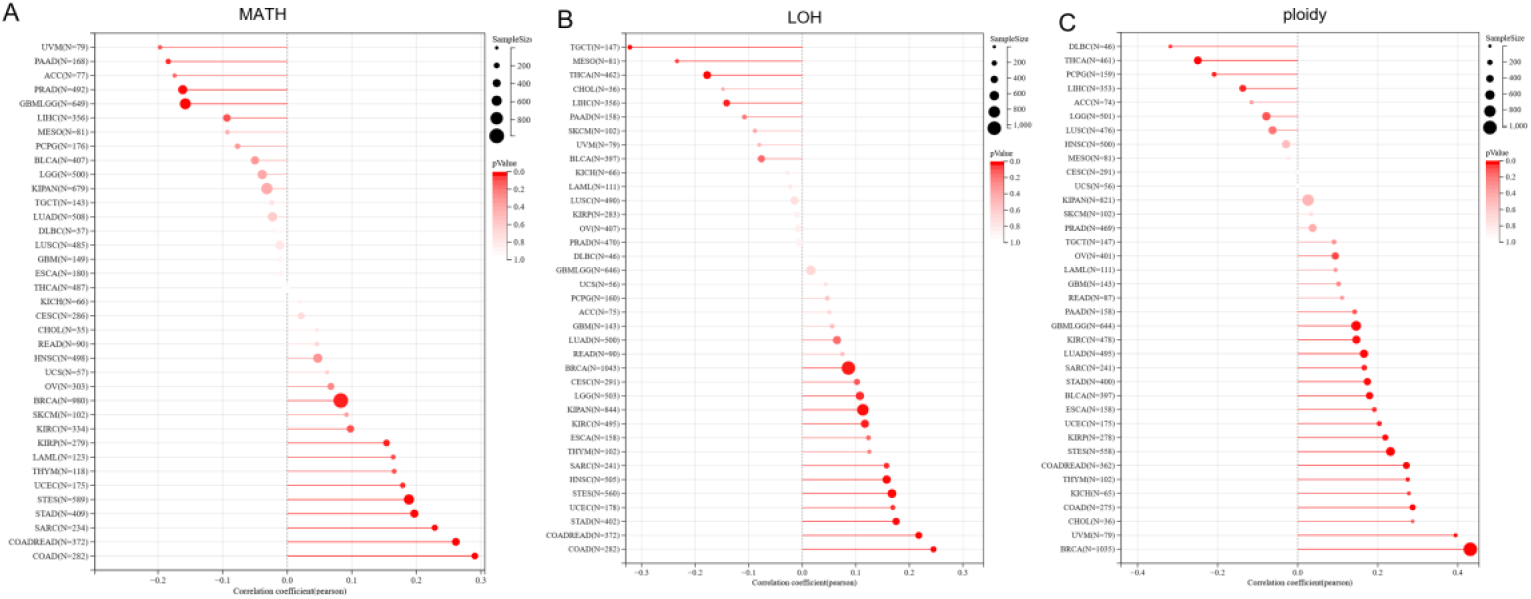
CEBPB expression correlates with genomic instability metrics across human cancers. (A) Pearson correlation coefficients between CEBPB expression and MATH scores across 33 TCGA cancer types, with STAD, KIRC, and LUAD showing significant positive associations. (B) Bubble plot illustrating CEBPB-LOH correlations across cancers, with bubble size representing -log10(p-value) and color indicating correlation strength (red: positive; blue: negative). (C) Chromosomal instability and aneuploidy (increased ploidy).

When examining these relationships specifically in STAD, CEBBP expression maintained significant positive correlations with MATH, ploidy, and LOH (all p < 0.01). These consistent associations across multiple genomic instability metrics suggest that CEBPB may function as a molecular amplifier of genomic chaos, potentially through regulating DNA damage response, chromosome segregation, or telomere maintenance pathways. The particularly strong association with LOH in gastric cancer implies that CEBPB might influence mechanisms of allelic imbalance and large-scale chromosomal alterations.

## Method and material

### Data Acquisition and Processing

The RNA-seq data from four Helicobacter pylori-infected gastric epithelial cell lines (AGS, GES-1, HGC-27, and MKN-45) were obtained from the Gene Expression Omnibus (GEO) database under accession number GSE295523. Raw sequencing reads were processed through a standardized bioinformatics pipeline. Quality control was performed using FastQC (v0.12.1), followed by adapter trimming with Trim Galore (v0.6.10) and alignment to the human reference genome (GRCh38) using STAR (v2.7.10a) [13,14]. Gene-level quantification was generated with featureCounts (v2.0.3) [15]. For gastric cancer clinical analysis, transcriptomic profiles and corresponding clinical data of stomach adenocarcinoma (STAD) patients were retrieved from The Cancer Genome Atlas (TCGA) database via the GDC Data Portal, encompassing RNA-seq data (FPKM normalized) and comprehensive survival information.

### Normalization of RNA-Sequencing Data

Comprehensive bioinformatics analysis was conducted using R software (version 4.4.1). Raw RNA-sequencing data were transformed into log_2_-based pseudocounts and filtered based on positive detection prior to quantile normalization performed via the preprocessCore

Bioconductor package (version 1.66.0). Subsequently, k-means clustering was implemented and batch effects were adjusted with the sva R package (version 3.52.0). Differential expression analysis was then executed on the normalized dataset using DESeq2, with significant differentially expressed genes defined as those exhibiting an absolute log_2_ fold change > 1 and an adjusted p-value < 0.01 [16].

### Functional analysis

Functional enrichment analysis of the identified gene modules was conducted via the R package clusterProfiler (version 4.12.6), which integrated resources from both the Gene Ontology (GO) Biological Process and the Kyoto Encyclopedia of Genes and Genomes (KEGG) pathway databases [17]. This integrative methodology supported a systematic annotation of the filtered differentially expressed genes (DEGs) and facilitated the detection of key signaling pathways significantly linked to immunotherapy resistance in Helicobacter pylori-associated gastric cancer.

### Survival and Prognostic Analysis

Survival analysis was performed to evaluate the prognostic significance of identified transcription factors in gastric cancer. Clinical data and corresponding RNA-seq expression profiles for stomach adenocarcinoma (STAD) were retrieved from The Cancer Genome Atlas (TCGA) database (https://portal.gdc.cancer.gov/).

The cohort was stratified into high- and low-expression groups for each transcription factor based on the median expression value. Kaplan – Meier survival curves were generated to compare overall survival (OS) between the two groups, and statistical significance was assessed using the log-rank test. Univariate Cox proportional hazards regression models were applied to calculate hazard ratios (HR) and 95% confidence intervals (CI) for each factor. Factors with a log-rank p-value < 0.05 were considered statistically significant. All survival analyses were conducted in R (version 4.4.1) using the survival (version 3.5-7) and survminer (version 0.4.9) packages. Data visualization was performed using ggplot2 (version 3.5.0).

### Immunological Profiling and Tumor Microenvironment Characterization

To delineate the immune landscape and investigate tumor microenvironment composition, we performed comprehensive immune infiltration analysis. The relative abundances of 28 distinct immune cell populations within each sample were quantified using the single-sample Gene Set Enrichment Analysis (ssGSEA) algorithm, implemented through the *GSVA* R package (version 1.48.3), based on well-curated gene signatures for immune cell types [18].

Subsequently, to deconvolve the transcriptomic data and estimate the specific proportions of 22 functionally defined immune cell subsets, we employed the CIBERSORT algorithm. This method utilizes support vector regression to infer cell-type composition from bulk tissue gene expression profiles [19]. Comparative analyses of immune infiltration scores were conducted between predefined sample groups (e.g., high vs. low CEBPB expression) using the Wilcoxon rank-sum test to identify statistically significant differences in immune cell abundance. Furthermore, Pearson or Spearman correlation analyses were performed to examine associations between the expression levels of target genes (e.g., *CEBPB*) and the infiltration levels of specific immune cell types, adjusting for potential confounding factors where appropriate. All analyses were executed in R software (version 4.4.1), with results visualized using the *ggplot2* and *pheatmap* packages.

## Discussion

Helicobacter pylori is a globally distributed pathogen and a well-defined gastric carcinogen. Chronic infection affects approximately half of the world’s population — with higher prevalence in low-income regions—and has been designated by WHO/IARC as a Group I carcinogen [20]. The bacterium drives chronic gastritis and secretes virulence factors such as CagA and VacA, which disrupt epithelial signaling and genomic integrity, initiating a pathological sequence from gastritis to atrophy, metaplasia, and ultimately gastric cancer [21]. Epidemiologically, H. pylori is the leading cause of gastric cancer worldwide; it is implicated in about 90% of non-cardia gastric tumors, and substantial evidence suggests that eradicating the infection could prevent a significant proportion of the global gastric cancer burden [22,23]. Therefore, our study is based on the established premise that H. pylori infection leaves a durable transcriptional and functional “imprint” on the gastric mucosa, promoting oncogenic processes—including chronic inflammation, reactive oxygen species (ROS) production, DNA damage, and immune suppression that collectively fuel tumorigenesis.

A central finding of our analysis is the identification of four ferroptosis-related genes, NOX4, MTCH1, GABARAPL2, and SLC2A3, whose expression is co-regulated by H. pylori status and predicts clinical outcome. Each gene has been implicated in cancer-relevant pathways, supporting their biological significance. NOX4, overexpressed in gastric tumors, drives ROS generation and modulates ferroptosis susceptibility [24]; its upregulation in H. pylori-positive tissues aligns with reports linking high NOX4 levels to poorer survival and increased tumor aggressiveness. MTCH1, a mitochondrial membrane protein, has recently been shown to promote ferroptosis through the FoxO1-GPX4 axis in cancer cells [25]. Although its role in gastric cancer remains less explored, MTCH1’s ability to induce lipid peroxidation and cell death in other malignancies suggests a plausible pro-ferroptotic function in H. pylori-driven carcinogenesis. SLC2A3 (GLUT3), a high-affinity glucose transporter, is markedly upregulated in gastric cancer and correlates with immune infiltration and adverse prognosis [26]. In gastric carcinoma, SLC2A3 enhances glycolytic flux and promotes M2-macrophage polarization, thereby reinforcing a pro-tumor microenvironment. Finally, GABARAPL2, an ATG8-family autophagy protein, recurs in gastric cancer prognostic signatures [27]. While its specific role in gastric cancer is not fully characterized, GABARAPL2 participates in autophagic transport and cellular stress responses. Notably, H. pylori infection is known to modulate autophagy in gastric epithelium, inducing LC3-positive autophagosomes and supporting the emergence of cancer stem-like traits [28]. Thus, the inclusion of GABARAPL2 in our signature resonates with evidence that H. pylori-driven autophagy contributes to tumor initiation.

Our integrated analysis highlights the dual impact of H. pylori on gastric carcinogenesis: it not only activates oncogenic networks within epithelial cells but also shapes an immunosuppressive tumor microenvironment. The co-differentially expressed genes identified in this study are enriched in oxidative-stress, apoptotic, and autophagy pathways—consistent with the known ability of H. pylori to induce ROS generation and DNA damage [29]. For instance, infection has been shown to perturb redox homeostasis through mechanisms such as the NOX-ROS-Nrf2/HO-1-ROS loop, thereby promoting ROS-mediated autophagy [30,31]. Moreover, H. pylori-induced autophagy in gastric epithelial cells further underscores the relevance of this process in infection-associated pathogenesis [32,33].

Mechanistically, CEBPB serves as a pivotal transcriptional hub whose dysregulated expression likely exerts direct control over downstream gene networks involved in cell cycle regulation and immune suppression. Through integrated transcriptomic analyses, this study systematically delineates, for the first time, the central role of CEBPB in remodeling the gastric tumor microenvironment induced by *H. pylori*. This transcription factor not only drives the malignant transformation of epithelial cells but also facilitates crosstalk between tumor cells and immune cells, thereby promoting the establishment of an immunosuppressive microenvironment. This finding provides a novel molecular perspective for understanding the pathogenesis of infection-associated gastric cancer and lays a theoretical foundation for developing therapeutic strategies targeting the CEBPB signaling pathway. By intervening in the CEBPB-mediated transcriptional network, it may be possible to achieve multi-level inhibition of *H. pylori*-related gastric carcinogenesis, underscoring the significant translational value of this work.

## Data availability

The datasets generated and/or analyzed during this study are available in the Gene Expression Omnibus (GEO) repository. All other data supporting the findings of this study are included in this published article.

## References

1. Rodriguez AM, Urrea DA, Prada CF. Helicobacter pylori virulence factors: relationship between genetic variability and phylogeographic origin. PeerJ. 2021;9:e12272. 10.7717/peerj.12272

2. Taddesse G, Habteselassie A, Desta K, Esayas S, Bane A. Association of dyspepsia symptoms and Helicobacter pylori infections in private higher clinic, Addis Ababa, Ethiopia. Ethiop Med J. 2011;49:109–16.

3. de Martel C, Georges D, Bray F, Ferlay J, Clifford GM. Global burden of cancer attributable to infections in 2018: a worldwide incidence analysis. Lancet Glob Health. 2020;8:e180–90. 10.1016/S2214-109X(19)30488-7

4. Kusters JG, van Vliet AHM, Kuipers EJ. Pathogenesis of Helicobacter pylori infection. Clin Microbiol Rev. 2006;19:449–90. 10.1128/CMR.00054-05

5. Zhang J, Wang W, Yan S, Li J, Wei H, Zhao W. CagA and VacA inhibit gastric mucosal epithelial cell autophagy and promote the progression of gastric precancerous lesions. Zhong Nan Da Xue Xue Bao Yi Xue Ban. 2022;47:942–51. 10.11817/j.issn.1672-7347.2022.210779

6. Sun J, Aoki K, Zheng J-X, Su B-Z, Ouyang X-H, Misumi J. Effect of NaCl and Helicobacter pylori vacuolating cytotoxin on cytokine expression and viability. World J Gastroenterol. 2006;12:2174–80. 10.3748/wjg.v12.i14.2174

7. Lin L, Xie B, Shi J, Zhou CM, Yi J, Chen J, et al. [IL-8 Links NF-κB and Wnt/β-Catenin Pathways in Persistent Inflammatory Response Induced by Chronic Helicobacter pylori Infection]. Mol Biol (Mosk). 2023;57:713–6.

8. Hu W, Chen ZM, Wang Y, Yang C, Wu ZY, You LJ, et al. Single-cell RNA sequencing dissects the immunosuppressive signatures in Helicobacter pylori-infected human gastric ecosystem. Nat Commun. 2025;16:3903. 10.1038/s41467-025-59339-4

9. O’Brien VP, Koehne AL, Dubrulle J, Rodriguez AE, Leverich CK, Kong VP, et al. Sustained Helicobacter pylori infection accelerates gastric dysplasia in a mouse model. Life Sci Alliance. 2021;4:e202000967. 10.26508/lsa.202000967

10. De Re V. Molecular Features Distinguish Gastric Cancer Subtypes. Int J Mol Sci. 2018;19:3121. 10.3390/ijms19103121

11. Kehrberg RJ, Bhyravbhatla N, Batra SK, Kumar S. Epigenetic regulation of cancer-associated fibroblast heterogeneity. Biochim Biophys Acta Rev Cancer. 2023;1878:188901. 10.1016/j.bbcan.2023.188901

12. Marusawa H, Chiba T. Helicobacter pylori-induced activation-induced cytidine deaminase expression and carcinogenesis. Curr Opin Immunol. 2010;22:442–7. 10.1016/j.coi.2010.06.001

13. Dobin A, Davis CA, Schlesinger F, Drenkow J, Zaleski C, Jha S, et al. STAR: ultrafast universal RNA-seq aligner. Bioinformatics. 2013;29:15–21. 10.1093/bioinformatics/bts635

14. Harrow J, Frankish A, Gonzalez JM, Tapanari E, Diekhans M, Kokocinski F, et al. GENCODE: the reference human genome annotation for The ENCODE Project. Genome Res. 2012;22:1760–74. 10.1101/gr.135350.111

15. Liao Y, Smyth GK, Shi W. featureCounts: an efficient general purpose program for assigning sequence reads to genomic features. Bioinformatics. 2014;30:923–30. 10.1093/bioinformatics/btt656

16. Love MI, Huber W, Anders S. Moderated estimation of fold change and dispersion for RNA-seq data with DESeq2. Genome Biol. 2014;15:550. 10.1186/s13059-014-0550-8

17. Yu G, Wang L-G, Han Y, He Q-Y. clusterProfiler: an R package for comparing biological themes among gene clusters. OMICS. 2012;16:284–7. 10.1089/omi.2011.0118

18. Hänzelmann S, Castelo R, Guinney J. GSVA: gene set variation analysis for microarray and RNA-seq data. BMC Bioinformatics. 2013;14:7. 10.1186/1471-2105-14-7

19. Newman AM, Liu CL, Green MR, Gentles AJ, Feng W, Xu Y, et al. Robust enumeration of cell subsets from tissue expression profiles. Nat Methods. 2015;12:453–7. 10.1038/nmeth.3337

20. Liu Y, Wei D, Deguchi Y, Xu W, Tian R, Liu F, et al. PPARδ dysregulation of CCL20/CCR6 axis promotes gastric adenocarcinoma carcinogenesis by remodeling gastric tumor microenvironment. Gastric Cancer. 2023;26:904–17. 10.1007/s10120-023-01418-w

21. Wang L, Gong W-H. Predictive model using four ferroptosis-related genes accurately predicts gastric cancer prognosis. World J Gastrointest Oncol. 2024;16:2018–37. 10.4251/wjgo.v16.i5.2018

22. Thrift AP, Wenker TN, El-Serag HB. Global burden of gastric cancer: epidemiological trends, risk factors, screening and prevention. Nat Rev Clin Oncol. 2023;20:338–49. 10.1038/s41571-023-00747-0

23. Tanaka C, Otani K, Tamoto M, Yoshida H, Nadatani Y, Ominami M, et al. Efficacy evaluation of upper gastrointestinal endoscopy screening for secondary prevention of gastric cancer using the standardized detection ratio during a medical check-up in Japan. J Clin Biochem Nutr. 2024;74:253–60. 10.3164/jcbn.24-28

24. Wang L, Gong W. NOX4 regulates gastric cancer cell invasion and proliferation by increasing ferroptosis sensitivity through regulating ROS. Int Immunopharmacol. 2024;132:112052. 10.1016/j.intimp.2024.112052

25. Wang X, Ji Y, Qi J, Zhou S, Wan S, Fan C, et al. Mitochondrial carrier 1 (MTCH1) governs ferroptosis by triggering the FoxO1-GPX4 axis-mediated retrograde signaling in cervical cancer cells. Cell Death Dis. 2023;14:508. 10.1038/s41419-023-06033-2

26. Lin L, Que R, Wang J, Zhu Y, Liu X, Xu R. Prognostic value of the ferroptosis-related gene SLC2A3 in gastric cancer and related immune mechanisms. Front Genet. 2022;13:919313. 10.3389/fgene.2022.919313

27. Qiu J, Sun M, Wang Y, Chen B. Identification and validation of an individualized autophagy-clinical prognostic index in gastric cancer patients. Cancer Cell Int. 2020;20:178. 10.1186/s12935-020-01267-y

28. Jia H, Wei J, Zheng W, Li Z. The dual role of autophagy in cancer stem cells: implications for tumor progression and therapy resistance. J Transl Med. 2025;23:583. 10.1186/s12967-025-06595-z

29. Shen X, Zhang W, Peng C, Yan J, Chen P, Jiang C, et al. In vitro anti-bacterial activity and network pharmacology analysis of Sanguisorba officinalis L. against Helicobacter pylori infection. Chin Med. 2021;16:33. 10.1186/s13020-021-00442-1

30. Lin Y, Liu K, Lu F, Zhai C, Cheng F. Programmed cell death in Helicobacter pylori infection and related gastric cancer. Front Cell Infect Microbiol. 2024;14:1416819. 10.3389/fcimb.2024.1416819

31. Li B, Lv X, Xu Z, He J, Liu S, Zhang X, et al. Helicobacter pylori infection induces autophagy via ILK regulation of NOXs-ROS-Nrf2/HO-1-ROS loop. World J Microbiol Biotechnol. 2023;39:284. 10.1007/s11274-023-03710-4

32. Olivera-Severo D, Uberti AF, Marques MS, Pinto MT, Gomez-Lazaro M, Figueiredo C, et al. A New Role for Helicobacter pylori Urease: Contributions to Angiogenesis. Front Microbiol. 2017;8:1883. 10.3389/fmicb.2017.01883

33. Donahue JP, Peek RM, Van Doorn LJ, Thompson SA, Xu Q, Blaser MJ, et al. Analysis of iceA1 transcription in Helicobacter pylori. Helicobacter. 2000;5:1–12. 10.1046/j.1523-5378.2000.00008.x

